# Which genetic variants in DNase I sensitive regions are functional?

**DOI:** 10.1101/007559

**Authors:** Gregory A. Moyerbrailean, Chris T. Harvey, Cynthia A. Kalita, Xiaoquan Wen, Francesca Luca, Roger Pique-Regi

## Abstract

Ongoing large experimental characterization is crucial to determine all regulatory sequences, yet we do not know which genetic variants in those regions are non-silent. Here, we present a novel analysis integrating sequence and DNase I footprinting data for 653 samples to predict the impact of a sequence change on transcription factor binding for a panel of 1,372 motifs. Most genetic variants in footprints (5,810,227) do not show evidence of allele-specific binding (ASB). In contrast, functional genetic variants predicted by our computational models are highly enriched for ASB (3,217 SNPs at 20% FDR). Comparing silent to functional non-coding genetic variants, the latter are 1.22-fold enriched for GWAS traits, have lower allele frequencies, and affect footprints more distal to promoters or active in fewer tissues. Finally, integration of the annotations into 18 GWAS meta-studies improves identification of likely causal SNPs and transcription factors relevant for complex traits.

## 1 Introduction

Genome-wide association studies (GWAS) have successfully identified large numbers of genetic variants associated with disease and normal trait variation (Mailman et al. 2007); yet a formidable challenge remains in determining the specific functional relevance of human DNA sequence variants. First, GWAS only identify large regions of association and in general, cannot directly pinpoint the true causative variants. Second, even after fine mapping, most of these genetic variants are located in non-coding regions that make it more difficult to infer the mechanism linking individual genetic variants with the disease trait. Third, we generally do not know in which cell-types/tissues, sequence variants identified by GWAS may have a functional impact.

Functional genomics data collected by ENCODE (The ENCODE Project Consortium 2012), Roadmap Epigenomics (Bernstein et al. 2010), and other groups (e.g., Visel et al. 2009) have provided a great deal of information on the tissue-specific regulatory regions of the human genome. At the same time, several computational methods have been developed that can exploit the correlation structure of these types of data to identify tissue-specific regulatory regions. For example, different types of histone modifications have been integrated using hidden Markov models (HMMs) (Ernst and Kellis 2010), and dynamic Bayesian networks (DBNs) (Hoffman et al. 2013). Open chromatin and sequence motifs have been integrated in different types of mixture models (CENTIPEDE (Pique-Regi et al. 2011), PIQ (Sherwood et al. 2014) and others (Boyle et al. 2012; Neph et al. 2012)). All these computationally and experimentally derived annotations for regulatory regions have been used to functionally characterize GWAS hits (Trynka et al. 2013; Schaub et al. 2012; Boyle et al. 2012; The ENCODE Project Consortium 2012; Neph et al. 2012; Ward and Kellis 2012). However, a simple positional overlap between a genetic variant and regulatory regions is not sufficient evidence to prove that the allelic status has a functional impact on binding. One of the major current limitations is that motif models for transcription factor (TF) binding are generally not sufficiently well calibrated to make a prediction of the sequence impact on binding.

Here, we have extended the CENTIPEDE approach to generate a catalog of regulatory sites and binding variants for more than 600 experimental samples from the ENCODE and Roadmap Epigenomics projects using recalibrated sequence motif models for more than 800 TFs. This is the most comprehensive catalog of regulatory variation annotated through integration of sequence and functional data to date. We then incorporated ASB information to provide additional empirical evidence and to validate the accuracy of the computational predictions. We also examined genomic properties of the annotations, identifying characteristics that separate variants that disrupt binding from those that do not, and demonstrated the action of natural selection on TF binding sites. Finally, we used our catalog to annotate and interpret variants associated with complex traits and we validate the allele-specific enhancer activity by reporter gene assay. Our results show that this strategy provides a general framework for the identification of regulatory variants and the determination of their functional role in complex traits.

## 2 Results

### 2.1 Identification of regulatory sequences and factor activity

The CENTIPEDE approach allows one to predict TF activity from integrating sequence motifs together with functional genomics data, and gains the most information from high-resolution data such as DNase-seq or ATAC-seq (Buenrostro et al. 2013). In the original CENTIPEDE approach, the sequence models are pre-determined; e.g, k-mers or previously defined position weight matrix (PWM) models. However, many motif models were created with very few sample sequences obtained from known TF binding sites and do not represent the full spectrum of sequence variation that can be tolerated without affecting binding. We have extended CENTIPEDE to readjust the sequence model for TF binding (Fig. 1A and Fig. S1) using DNase-seq data and sequence orthologs (Methods). Compared to the original motif models the consensus sequence is largely maintained in the recalibrated motifs (Fig. S6). When we consider ChIP-seq peaks as validation we obtain superior precision recall characteristics (Supplemental S6.1 and Fig. S7) and a much higher correlation with the prior probability of binding calculated by CENTIPEDE (Suplemental S6.2 and Fig. S8)

**Figure 1:**
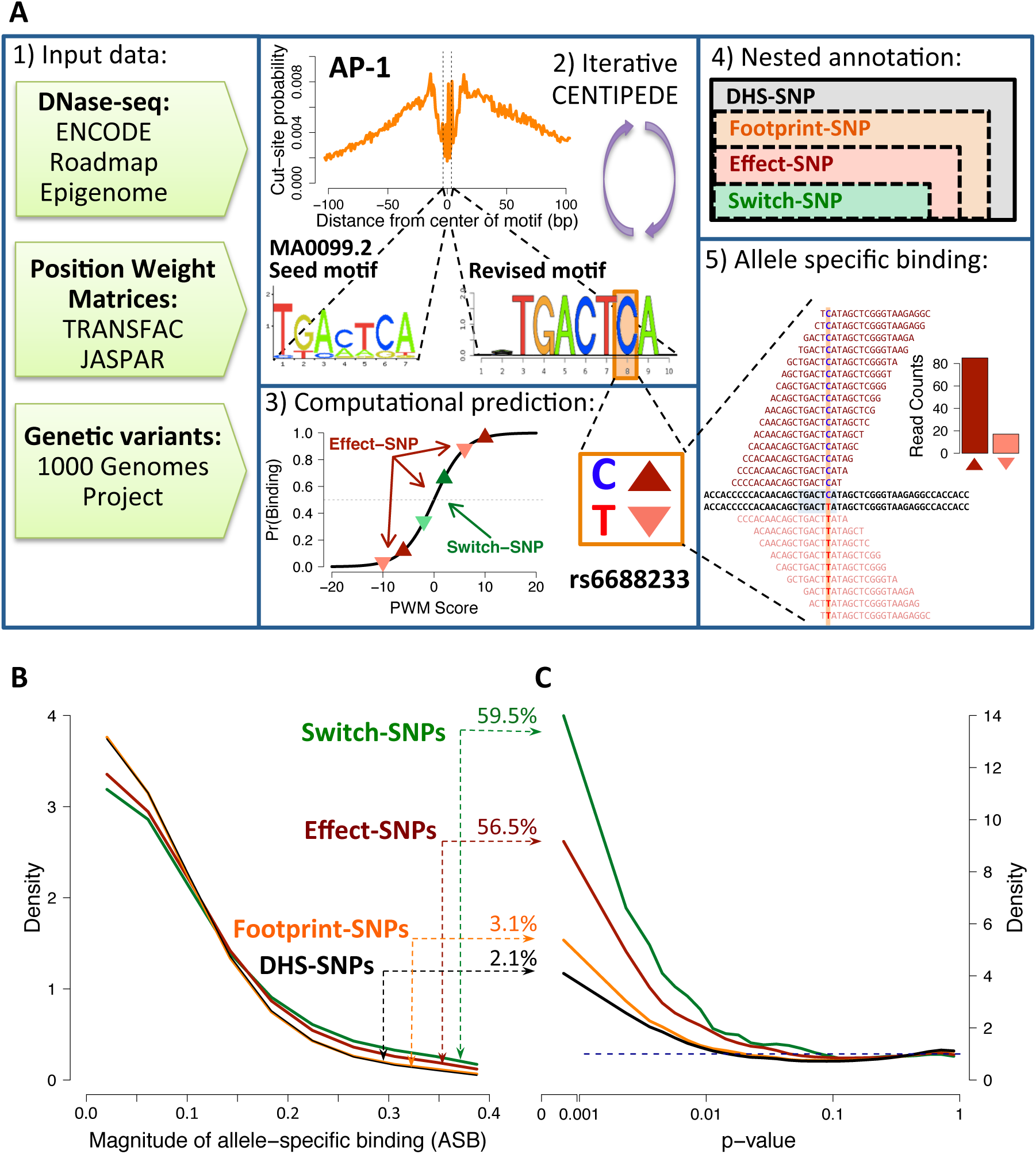
**Identification of allele-specific binding.** (A) A visual description of the methods. [(A1) Data sources. (A2) Iterative process of using CENTIPEDE to revise sequence models and call footprints. (A3) Computational predictions of factor binding. SNPs are categorized by the degree to which they are predicted to impact binding. (A4) The nested categories of functional annotations. (A5) Use of allele-specific analysis to detect differential binding.] (B) SNPs predicted to have effects on binding are enriched for a large magnitude of allelic imbalance |(allele ratio - 0.5)|. Each line represents a density plot of the magnitude for SNPs within a functional category. Numbers on arrows represent the expected proportion of true signal (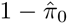) for each category. (C) ASB p-value densities for different SNP categories (the dotted blue line represents the null distribution).

Across all 653 tissues, we identified a total of 6,993,953 non-overlapping footprints corresponding to 1,372 active motifs and spanning 4.15% of the genome. Each individual sample contained, on average, 280,000 non-overlapping footprints for 600 motifs and spanning 0.162% of the genome, indicating that footprints are highly tissue specific. Considering all SNPs from 1000 Genomes Project (1KG) at any allele frequency (even singletons), we found 5,810,227 unique genetic variants in active footprints (footprint-SNPs), 3,831,862 of which are predicted to alter the prior odds of binding ≥20-fold (effect-SNPs) based on the logistic sequence model hyperprior in the CENTIPEDE model (Fig. 1A3, Supplemental eq. 2). In some cases, effect-SNPs increase (or decrease) the binding affinity of a sequence with an already high (or low) affinity, and for these cases the functional impact is likely to be small. However, 264,965 of these SNPs are predicted to have a larger effect on the binding of a factor (switch-SNPs), switching from a high prior probability of binding (>0.5) to a low probability (<0.5).

### 2.2 Analysis of allele-specific binding using DNase-seq data

To validate the SNP functional predictions, we used our recently developed approach, Quantitative Allele-Specific Analysis of Reads (QuASAR) (Harvey et al. 2014) to perform joint genotyping and ASB analysis within DNase I hypersensitivity (DHS) regions. The parameters of the QuASAR model also allowed us to detect tissues with chromosomal abnormalities or samples from pooled individuals (Supplemental S4.2), which we excluded from ASB analysis (Table S5; Fig. S9 and S10). Across the remaining 316 samples, we identified 204,757 heterozygous SNPs in DHS sites (DHS-SNPs) with coverage >10x. Based on our classification, among DHS-SNPs: 55,044 are footprint-SNPs, 26,773 are effect-SNPs, and 5,991 are switch-SNPs. Overall, the predictions are highly concordant with the direction of ASB; 88% of the sequence models show positive correlation between the predicted and observed ASB (Supplemental S5.2, Fig. S11). Each of the nested SNP functional categories have marked differences in p-value distribution (Fig. 1C) for the QuASAR test of ASB. Compared to what would be expected from the null uniform distribution, effect-SNPs and switch-SNPs have 8x and 14x times more SNPs with *p* < 0.001 respectively, showing that our functional annotations can predict ASB. Furthermore, these enrichments for lower p-values are much higher than those of DHS-SNPs (4x) and footprint-SNPs (6x), indicating that identifying SNPs in DHS regions and/or footprints alone is not enough to predict functional effects.

We then examined the observed allele ratios (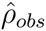) across different CENTIPEDE annotations. Fig. 1B shows that effect-SNPs and switch-SNPs tend to have a higher magnitude of allelic imbalance. If we consider SNPs that are in the tail of the distribution 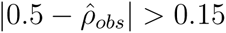, effect-SNPs show a moderate but significant enrichment (1.29-fold, Fisher p = 1.1×10^−229^) compared to DHS-SNPs.

All these results indicate that the following three categories of SNPs: effect-SNPs, footprint-SNPs that are not effect-SNPs, and the remaining DHS-SNPs, have marked differences in their probability to show evidence for ASB (Table 1). We therefore followed the strategy of Benjamini and Bogomolov (2014) to perform multiple testing correction in each category separately using Storey’s procedure (Storey 2003). At an FDR threshold of 20%, we detect 3,217 unique loci displaying significant ASB (Table 1), hereafter referred to as ASB-SNPs. Several of the ASB-SNPs were significant in more than one cell-type, giving a total of 4,940 observations of ASB-SNPs. Based on the proportion of true null hypothesis (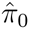) estimated by the Storey FDR procedure, we estimate that 56% of the effect-SNPs show evidence of ASB. While this conservative estimate can be considered a lower bound, it is still much higher than the estimates for DHS-SNPs (2.1%) and footprint-SNPs (3.1%), indicating that most SNPs in DHS regions and even in the putative binding sites do not affect binding.

**Table 1:** **Summary of Allele-Specific Binding SNPs.** Each row represents a category that is a subset of the category in the previous row. Each column reports the number of heterozygous SNPs, SNPs displaying significant ASB (20% FDR), and the estimated proportion of non-null hypotheses using Storey’s q-value approach. In parentheses are reported the numbers for SNPs that are not present in any of the subsequent subsets and are the basis for our partitioned q-value approach to detect ASB-SNPs.

| | # Het SNPs | # ASB (20% FDR) | $1 - \hat{\pi}_0$ |
| --- | --- | --- | --- |
| All DHS Hets | 204,757 (179,137) | 0 (0) | 2.1 (1.7)% |
| Footprint-SNPs | 55,044 (42,098) | 0 (0) | 3.1 (0.3)% |
| Effect-SNPs | 26,773 (26,773) | 3,217 (3,217) | 56.5 (56.5)% |

### 2.3 Characterization of effect-SNPs

Regions of the genome with demonstrated molecular function (e.g. genic regions) generally show reduced diversity (McVicker et al. 2009) and a site frequency spectrum skewed towards rare variants. This is due to negative (purifying) selection, which prevents deleterious alleles from reaching high frequencies in the population. We investigated whether a similar skew in the site frequency spectrum exists at functional non-coding variants (effect-SNPs). We observed that effect-SNPs are slightly more likely to be rare compared to those that do not affect binding (Fig. 2A), displaying a mild but significant 1.08-fold enrichment for having a MAF <1%. This observation is analogous to what has been seen among coding variation, where rare (< 0.5%) variants are 1 to 2 times more likely to be non-synonymous changes than synonymous (The 1000 Genomes Project Consortium 2012). Of effect-SNPs with MAF < 1%, a slightly higher proportion are predicted to increase binding of a factor (54%), rather than decrease binding (46%) relative to the major allele.

**Figure 2:**
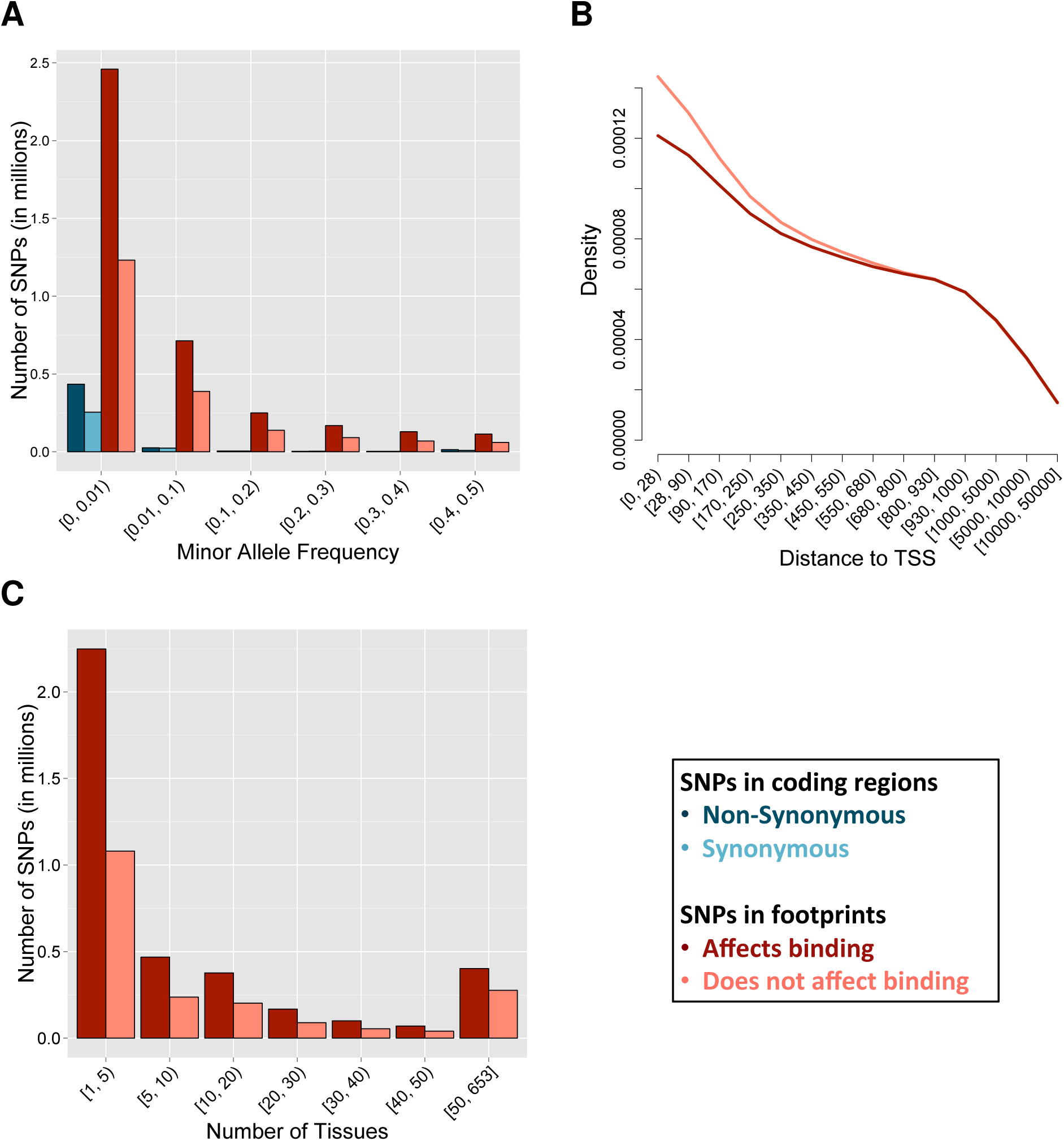
**Characterization of binding SNPs.** (A) Comparison of the minor allele frequency of SNPs predicted to affect binding (dark red) or not (light red). Minor allele frequency at coding SNPs, separated into non-synonymous (dark blue) and synonymous (blue), is shown for comparison. MAF is in bins of 10%, with the exception of rare (MAF < 1%) SNPs. (B) Proportion of SNPs at increasing distance from the nearest transcription start site (TSS) up to 50Kb. Fewer SNPs affecting binding (dark red) are present near a TSS compared to those with no effect (light red). Distance is absolute distance, regardless of direction (up- or downstream) from TSS. (C) Stratification of footprint-SNPs by the number of tissues for which the footprint was predicted active, colored by binding effect (dark red) or not (light red). Number of tissues is binned by 5 or 10 until 50, where the remainder is binned.

eQTL studies have found that variants associated with gene expression tend to occur close to the transcription start site (TSS) (Veyrieras et al. 2008; Gaffney et al. 2012; McVicker et al. 2013). We detect a similar trend among our annotations, with 83% of footprint-SNPs occurring within 100kb of the TSS. However, we find a 1.12-fold depletion of effect-SNPs within 300 bases of a TSS (Fig. 2B), probably because we are focusing on the core promoter region (Cooper et al. 2006). We also discovered a 1.18-fold enrichment for effect-SNPs in motifs active in 5 or fewer samples (Fig. 2C). These results support the hypothesis that factors binding closer to the TSS and/or active in many tissues are housekeeping factors and those that recruit the transcriptional machinery, and therefore changes in binding affinity are more likely detrimental to the cell.

Since allele frequency can be correlated with distance to the TSS, and shared footprints may also be more common at the promoter region, we tested all these features together in a joint model to control for one variable being a confounder of another (Methods). All three factors are significant predictors when considered together in a multiple regression logistic model, and the direction of the effect is the same as when considered separately (Table S6).

### 2.4 Motif-wise characteristics of regulatory variants

To examine the distribution of ASB-SNPs across the different regulatory factors, we calculated the ASB enrichment ratio (Fig. S12, Supplemental S7.3). At a nominal p-value cutoff of *p* < 0.01, we detect 32 motifs enriched for ASB and 56 depleted for ASB (Fig. 3A; Table S8). In cases where multiple motifs correspond to a single factor, we asked if all binding sites were similarly enriched or depleted for ASB-SNPs. We generally found this to be the case (Table S9), most notably for the factor AP-1. We see the same pattern for motifs significantly depleted of ASB-SNPs, such as CTCF and E2F (Table S9). ASB enrichment ratios are also consistent across factors with similar functions. For example, three factors in addition to AP1 with roles in the immune response, CREB (Zhao and Brinton 2004), c/EBP (Hu et al. 2000), and NF-*κ*B (Thomas et al. 1997) are over 2-fold enriched for ASB-SNPs within their binding sites (Table S10).

**Figure 3:**
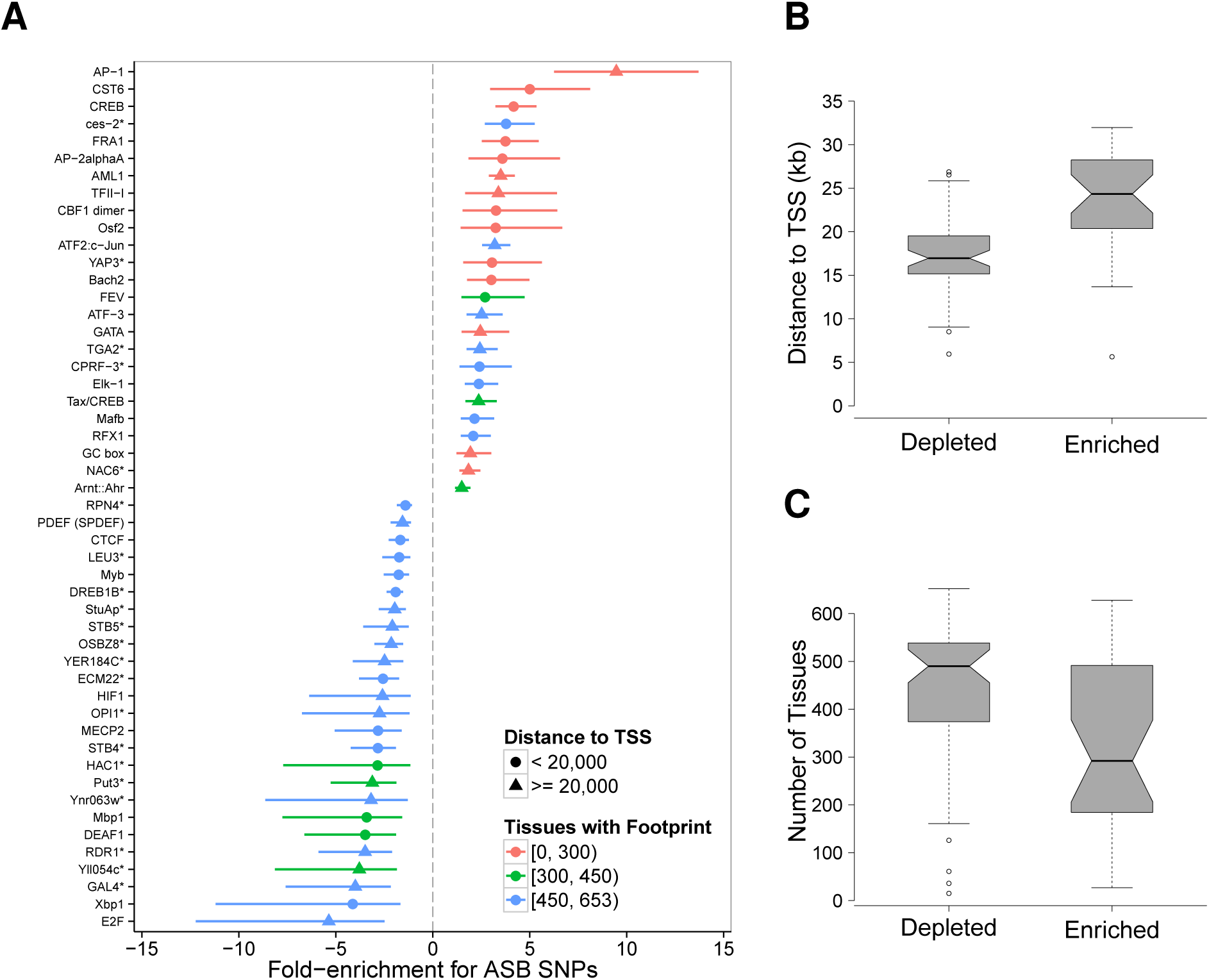
**Characterization of binding elements.** (A) Plot showing factors whose binding sites are significantly enriched or depleted for ASB variants (p-value <0.01), with indication of the number of tissues affected (color) and the median distance to the TSS (shape). Horizontal lines represent the 95% confidence interval of the ASB enrichment ratio. An asterisk denotes a possible human analog for the specified factor. Redundant motifs were excluded from the plot. (B) Barplot showing the distance to the nearest TSS between motifs either enriched or depleted for ASB-SNPs. (C) Barplot showing the number of tissues where a motif was predicted to be active for motifs either enriched or depleted for ASB-SNPs.

We then examined the genomic characteristics at motif instances to identify features that distinguish motifs enriched for ASB versus those that are not. We found that motifs enriched for ASB were significantly farther from the TSS, having an average median distance to the TSS of 23kb compared to 17kb for those depleted (*t*-test *p* = 2.8×10^−7^; Fig. 3B). Furthermore, motifs enriched for ASB were active in significantly fewer samples, on average active in 20% vs 40% for those depleted (*t*-test *p* = 5.2×10^−9^; Fig. 3C), indicating that TF with motifs with a high degree of ASB effects tend to be active in fewer tissues.

### 2.5 Evidence for motif-wise selection in TF binding sites

An important question in evolutionary biology is the extent to which selection has acted on *cis*-regulatory elements in humans (Wray 2007). While methods are being developed to address this question (Arbiza et al. 2013; Smith et al. 2013), such methods have only been applied to a narrow subset of TFs, and, in the case of Smith et al. (2013), rely on RNA expression data to classify mutations as up- or downregulating transcription relative to the reference enhancer sequence. Given our categorization of footprint-SNPs relative to their effect on factor binding, we performed an initial survey of selection across TF binding sites using a test similar to the McDonald-Kreitman (MK) test (McDonald and Kreitman 1991) (see Supplemental S7.4, Fig. S3). Applying our modified MK test, we obtained a selection score for factor motifs with sufficient sites in each category (Fig. 4, Table S11). At an FDR of 1%, we observe 84 factors whose binding sites are enriched for fixed functional differences (higher selection scores), suggestive of positive selection acting on those sites. Among the top scoring motifs are several factors that regulate neural and developmental processes, including POU1F1, PHOX2B, DBX2, UNCX, and YY1. Among the factors with the lowest selection scores, suggestive of purifying selection, we find ARNT, RBPJ, CREB1, POU2F2, and MYC.

**Figure 4:**
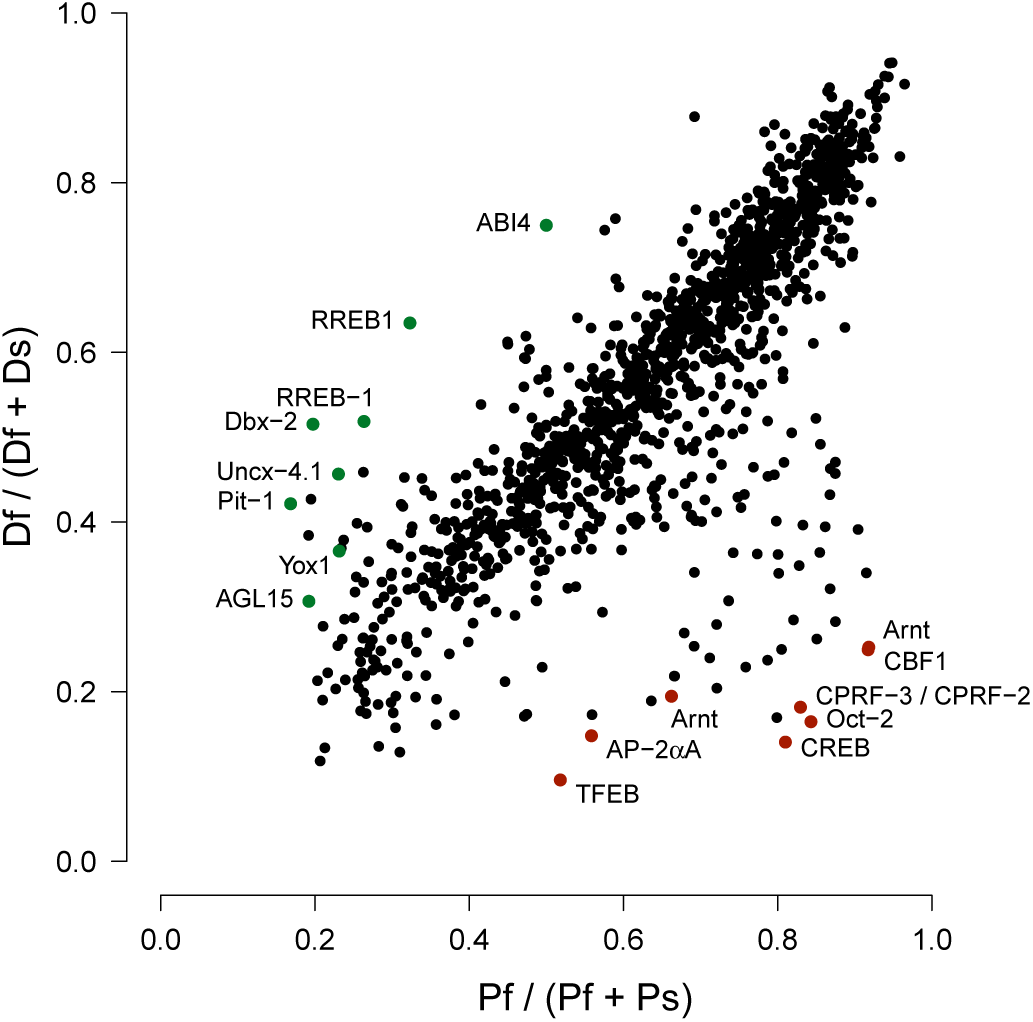
**Examining selection on TF binding sites.** Comparison of fixed functional (D*_f_*) to fixed silent (D*_s_*) (y-axis) versus polymorphic functional (P*_f_*) to polymorphic silent (P*_s_*) (x-axis) variants across all of the binding sites for each TF examined. Scores towards the top left are suggestive of positive selection (excess of fixed functional changes), with several of the top significant examples indicated by green points and the factor name. Scores towards the bottom right are suggestive of negative (purifying) selection, with several of the top significant examples indicated by red points and the factor name.

As a majority (994) of the factors showed mild but significant evidence for purifying selection, we next asked whether these sites were influenced by background selection from nearby genes (Fig. S15). We find a mild but significant positive correlation between selection score and median TSS distance (Spearman *ρ* = 0.16, *p* = 5.6×10^−9^). Additionally, there is a negative correlation between tissue specificity and selection score (Spearman *ρ* = -0.20, *p* = 1.2×10^−13^). While some of the selection signal may come from nearby genes under selection, there does appear to be a pattern of selective constraint on broadly active factors binding in promoter regions.

**Figure 5:**
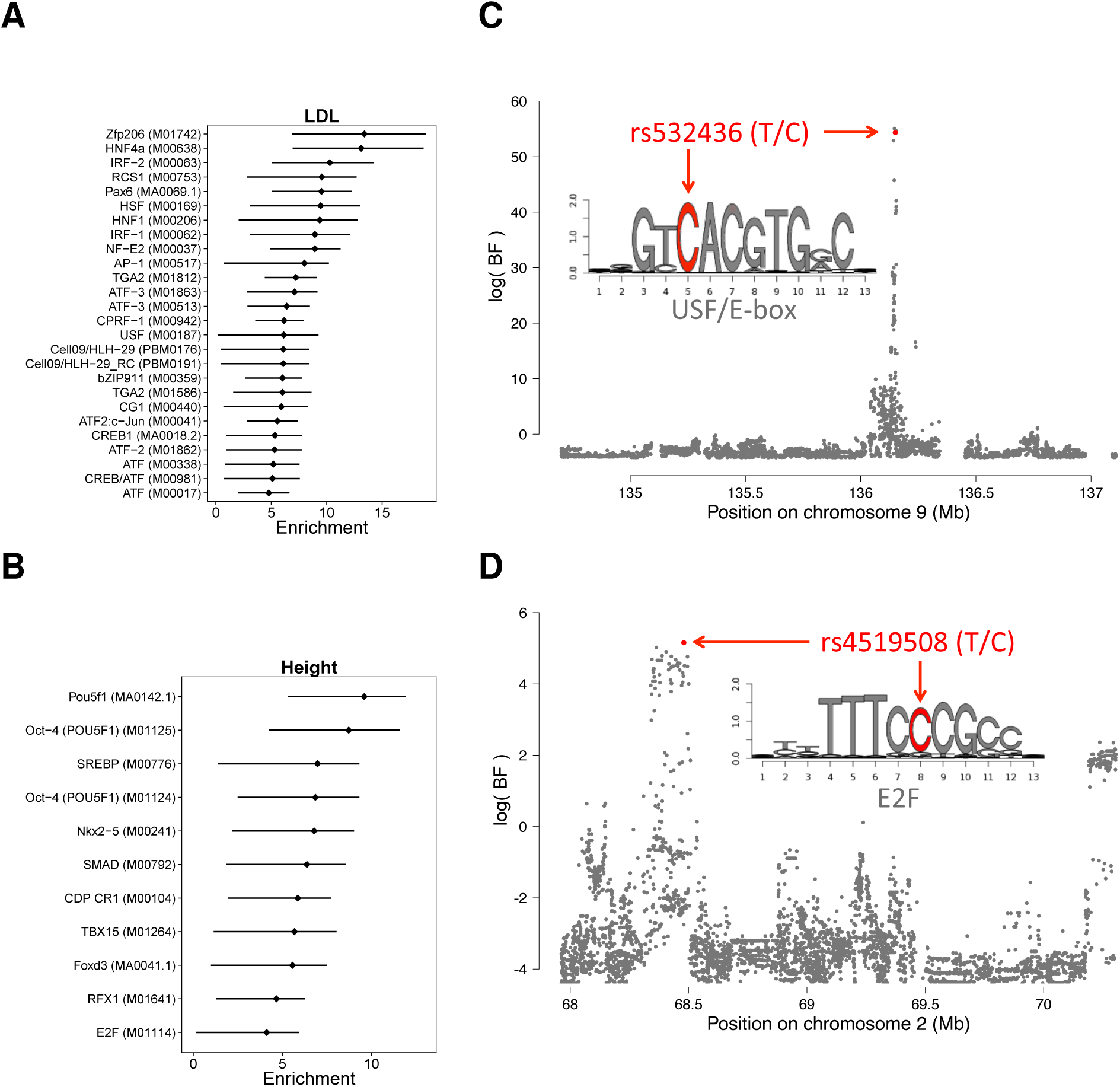
**Integration of annotations into GWAS results.** (A/B) Enrichment (*log*_2_(change in prior odds w.r.t the baseline model)) of factors for association with (A) low-density lipoprotein levels and (B) height. Error bars are drawn for 95% confidence intervals. (C/D) Association plots showing the Bayes factor of each SNP in the displayed region for (A) low-density lipoprotein levels and (B) height. Shown in red are SNPs with a posterior probability of association >0.4.

### 2.6 Enrichment of binding variants in GWAS data

Given that our annotations comprise predicted and observed functional effects across multiple cell-types/tissues, we asked if they could help interpret genomic hits reported in the GWAS catalog. To account for the possibility that the reported SNP is not itself causal, we defined the GWAS hit regions to include SNPs in linkage disequilibrium using an *r*^2^ cutoff of 0.8 (from 1KG Project). In GWAS hit regions, we compared the proportion of effect-SNPs and ASB-SNPs to footprint-SNPs and found a 1.11-fold enrichment for effect-SNPs and a 1.22-fold enrichment for ASB-SNPs (*p* < 2.2 × 10^−16^, 95% CI: 1.10 - 1.14 and *p* < 5 × 10^−3^, 95% CI: 1.06 - 1.39, respectively).

We next asked whether ASB sites may be used to improve existing functional annotations. Adding our annotation to category 2 SNPs from the RegulomeDB (Boyle et al. 2012) (SNPs with multiple regulatory annotations, but not shown to be functional), we detect a 1.6-fold enrichment for GWAS hits compared to category 2 SNPs alone (*p* = 6.11×10^−5^, 95% CI: 1.27 - 1.99), further highlighting the importance of identifying functional sites at a base-pair resolution, as a large number of genetic variants in regulatory sites are not functional.

To test if the annotated effect-SNPs can give a mechanistic support for SNPs associated with complex traits, we integrated them into GWAS meta analyses for 18 traits (see Table S14) using the recently developed hierarchical model fgwas (Pickrell 2014). For each trait, we identified factors whose binding sites were enriched for associated SNPs (Fig. 5, Fig. S13 and Table S7). Overall, we observed enrichment for biologically relevant factors; for example, factors enriched for height-associated SNPs include the embryonic stem cell factor POU5F1, consistent with previous observations of genetic variants associated with height being enriched in embryonic stem cell DHS sites (Trynka et al. 2014). We also see enrichment for the developmental regulators TBX15, FOXD3, and NKX2-5. From a study of low-density lipoprotein (LDL) levels in the blood, enriched factors include the liver-specific factor HNF4A, as well as several regulators of immune function, including SPIB, CREB1, IRF1, IRF2, and NR3C1 (GR).

Our high resolution annotations allowed us to dissect the most likely functional variant (posterior probability of association, PPA > 0.2) in 88 previously identified GWAS regions (Fig. S16, Table S13). We performed reporter gene assays for six SNPs to validate the predicted allelic effects on gene expression and the underlying molecular mechanism (Methods, Table S15). Among the regions tested we validated that five have enhancer/repressor activity and four have variants with allele-specific activity. These include rs4519508 (*p* = 0.041) and rs4973431 (*p* = 0.035) associated with height, rs532436 (*p* = 0.048) associated with blood LDL levels and rs6730558 (*p* = 0.022) associated with mean red cell volume.

rs4519508, associated with a 2.1cm decrease in height (Lango Allen et al. 2010), is in a binding site for the cell-cycle regulator family E2F (Fig. 5D). Our annotation increased the PPA from a baseline of 10.5% to 44.4%, and it is the highest associated SNP in the association block (Fig. S14A). We found that this E2F footprint is active in >300 tissues (most of them fetal) and we detected ASB at this SNP in lung fibroblasts, validating that the reference allele at rs4519508 confers stronger binding than the alternate. Interestingly, in the reporter assay we observed 1.3-fold increased expression in the presence of the alternate allele, suggesting that at this location, E2F is acting as a repressor. Finally, this SNP is located within the promoter of PPP3R1, a regulatory subunit of calcineurin important for cardiac and skeletal muscle phenotypes; and a SNP in the same region has been shown to be associated with endurance (He et al. 2010) in humans. The p-value of association for this GWAS locus (*p* = 8.1 × 10^−6^) does not reach genome-wide significance in the height meta-analysis data we used (Lango Allen et al. 2010). However, in a recent more extensive meta-analysis for height (Wood et al. 2014) this locus achieves genome-wide significance *p* = 8.4 × 10^−10^, demonstrating that our annotation can be useful to rescue relevant loci.

Finally, a SNP associated with LDL levels, rs532436, is within a footprint for USF, an E-box motif (Fig. 5C). Adding our annotation increased the PPA of the SNP from 39.7% to 94.7% (Fig. S14B). We found that the alternate allele, associated with a 0.0785 mg/dL increase of LDL in the blood, is predicted to have a lower binding probability and results in 2.5-fold lower expression, compared to the reference allele. This SNP is identified by GTEx as an eQTL for two proximal genes in whole blood: ABO (*p* = 5 × 10^−5^) and SLC2A6 (GLUT6, a class III glucose transport protein; *p* = 8 × 10^−5^). The SNP has an opposite effect on expression of the two genes, with the alternate allele showing lower expression for ABO and higher expression for SLC2A6.

These results show that our integrated analysis provides support for likely mechanisms linking regulatory sequence changes to complex organismal phenotypes. Furthermore, these mechanisms can be directly investigated through molecular studies, providing additional support that these sequence changes are truly functional.

## 3 Discussion

We have developed an approach for assessing functional significance of non-coding genetic variants. Our strategy integrates sequence information with functional genomics data to precisely annotate TF binding, and predict the impact of single nucleotide changes on these regions. By borrowing data from ENCODE and Roadmap Epigenomics, we generated one of the most comprehensive catalogs available to date annotating regulatory regions and functional genetic variants across the genome. We found that genetic variants that impact TF binding are depleted in the core promoter regions, tend to have low allele frequency and are enriched in tissue-specific footprints. These properties largely reflect the family-wise characteristics of motifs, which are further reflected in signals of selection. Finally, we showed how regulatory annotations improve the identification of potential causal SNPs in GWAS, and we provide examples of molecular mechanisms behind the association signals for height and blood lipid levels.

Thus far the most common approach to make use of functional genomics data to identify regulatory variants, assumes that each SNP in a regulatory region is equally likely to be functional. A key finding in this study is that genetic variants in regulatory active sequences, as defined by DNase I sensitivity and footprinting, are in very large proportion silent; only 2.1% of SNPs in DHS regions and 3.1% of SNPs in CENTIPEDE footprints are estimated to have ASB. This is analogous to SNPs in coding regions, where most genetic changes are synonymous and do not result in an amino acid change (The 1000 Genomes Project Consortium 2012). The sequence model developed in this study provides a very useful filter for non-coding genetic variants that are not functional, resulting in a tissue-specific and motif-specific annotation of effect-SNPs (56.5% of which are estimated to have an impact on ASB). This is crucial information to take into account when we attempt to understand the molecular mechanism behind GWAS hits and evolutionary signals of selection. As additional functional genomics studies are performed, across larger sample sizes, tissue types and cellular conditions, it will be important to further determine the functional subset of regulatory variants within binding sites to achieve greater power in functionally annotating genetic variants associated with complex traits.

A key feature of our annotation is that it spans a large collection of tissues and transcription factor motifs. This allowed us to trace some of the evolutionary history of TF binding and identify evolutionary constraints on specific molecular functions, which may reflect selective pressures during human history. For example, we observed that immune TFs are enriched for ASB sites, which supports the hypothesis that this may be a consequence of human adaptations to pathogen exposures (Miller et al. 1976). On the other hand, we identified neural development TFs that may have undergone positive selection in humans. The large number of regulatory variants predicted in our study, together with previously reported eQTL signals, and the overall relevance that they have in explaining complex traits provides further support for polygenic models of complex traits in humans. By taking advantage of the factor-specific annotations in our study, we identified motifs that are enriched for regulatory variants associated with relevant GWAS traits; e.g., immune TFs in the lipids study, and developmental TFs for height. Overall, the GWAS meta-analysis and selection signals in our study support the concept that variation in binding sites has been a major target of evolutionary forces and contribute to disease risk and complex phenotypes in human populations.

## Methods

### Identification of active regulatory sites and motif recalibration

We used 1,949 PWM sequence models from the TRANSFAC (Matys et al. 2006) and JASPAR (Sandelin et al. 2004) databases to scan the genome for a set of representative motif matches (Supplemental S3.1). For each sequence model, we used the sequences to calculate a new PWM model which we then used to scan the genome and identify all genome-wide motif matches using a two step approach:

*Step 1: Initial CENTIPEDE scan and motif recalibration.* After scanning the genome for motif matches (using 1949 seed motifs), we extracted DNase-seq data at these sites using 653 samples publicly available from the ENCODE and Roadmap Epigenomics projects (Supplemental S1 and S2.1). The motifs and samples used are summarized in Tables S1 and S2. For each motif and only for this initial step, we used a reduced subset of motif matches that include the top 5,000 instances on the human genome and up to 10,000 additional low-scoring sequences (Supplemental S3.1). These low scoring motif instances are human sequences that have orthologous high scoring motif instances in the chimp or rhesus genome. We then applied the CENTIPEDE model to survey TF activity for each 1,272,697 tissue-TF pair. For each pair we then determined that the TF is active if the motifs instances that exhibit a CENTIPEDE footprint can be predicted from the PWM score (Z-score > 5, Fig. S4 and S5). Using this criterion, we determined that 1,891 TF motifs are active in at least one tissue. The full list of motifs active in each tissue can be found in Table S12. We then recalibrated the PWM model for each active motif using the sequences of all motif matches that have a DNase-seq footprint (CENTIPEDE posterior >0.99).

*Step 2: Full genome CENTIPEDE scan and genetic variant analysis.* Using these newly updated sequence models we scanned the human genome for all possible matches both to the reference and to alternate alleles from genetic variants catalogued in the 1KG Project (The 1000 Genomes Project Consortium 2012) and used the CENTIPEDE algorithm to assess the probability that each motif instance is bound by a TF. In this second step, we included all high and low scoring PWM matches down to a CENTIPEDE prior probability of binding of 10% (Supplemental eq. 2 and S3.2).

### ChIP-seq validation of the revised sequence motif models

To evaluate whether the updated sequence models derived from DNase-seq data are better at predicting TF binding than the original seed motifs, we compared to ChIP-seq data available for a small set of TFs from the ENCODE project (as these data are generated in independent experimental assays that should be highly TF-specific). Using precision recall operating characteristic (P-ROC) curve analysis (see Supplemental S6.1), we determined that for a given precision (precision = 1 - FDR, false discovery rate), the updated sequence models have higher recall (sensitivity) than the original PWM in detecting ChIP-seq peaks (Fig. S7). Additionally, we compared the correlation between the prior probability of binding (calculated by CENTIPEDE based on the PWMs) and the number of ChIP-seq reads overlapping motif matches (Supplemental S6.2).

### Identification of allele-specific binding

Starting from raw sequencing reads, we used a custom mapper (Degner et al. 2012) to align the reads to the hg19 reference genome. As allelespecific analysis is extremely sensitive to mapping errors and PCR duplicates, we employed several methods to reduce these sources of potential bias (Supplemental S2.2 - S2.4). To detect allele-specific binding, we applied QuASAR (Harvey et al. 2014) to the processed read data. QuASAR first genotypes SNPs in the dataset using the read counts, then determines the likelihood of allelic imbalance at each heterozygous site. To adjust for multiple testing, we used the *q*-value method (Storey 2003) on the *p*-values produced by QuASAR.

### Annotation of ASB with binding predictions

To determine which SNPs displaying ASB fall within a predicted footprint, we overlapped DHS-SNPs identified by QuASAR with CENTIPEDE footprints for each sample. We classified a SNP as having an effect on binding if the difference in the prior log odds ratio (from the logistic sequence model in CENTIPEDE, Supplemental eq. 2) between the two alleles was ≥3, indicating a ≥20-fold change in the prior odds of TF binding. To generate a final set of annotated SNPs, we aggregated the data from each sample and motif into one table. For cases where a SNP is within multiple predicted binding sites, we selected the factor whose CENTIPEDE likelihood ratio was the greatest. SNPs were then partitioned based on their predicted effect on binding into three non-overlapping categories: 1) SNPs in predicted footprints whose binding effect is in the direction predicted, 2) all other SNPs in footprints, 3) all other DHS-SNPs. Because each annotation has a different prior expectation of being functional, we readjusted for multiple testing within each annotation separately using the *q*-value method (Storey 2003) on *p*-values produced by the QuASAR model.

### Regression model for binding effect

To see which features of a SNP were predictors of functional effect, we performed multiple regression analysis using a logistic model considering the dependent binary variable *E_l_*, indicating whether the footprint-SNP, *l*, is also an effect-SNP.

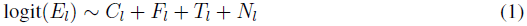

We considered the following variables related to the probability of a footprint-SNP being an effect-SNP: the log fold-change in factor affinity predicted by the sequence model (*C_l_*); the minor allele frequency (*F_l_*); the absolute distance to the nearest transcription start site (*T_l_*); the number of tissues for which the motif containing the footprint-SNP was predicted to be bound (*N_l_*). The model was fit using the GLM function in R.

### Identification of selection signals on TF motifs

To identify divergent TF binding sites, we used the UCSC liftOver tool on binding sites without a polymorphism to obtain orthologous regions in the chimpanzee genome. Using the PWM model, we calculated PWM scores on the chimpanzee sequences. Sites where the prior probability of binding differ from the humans sites were classified as divergent, and were further categorized by the difference in binding affinity: “functional” for sites that change ≥20-fold between species (analogous to effect-SNPs), and “silent” for those that do not. For the binding sites containing a polymorphism, we separated them into similar groupings where the change is functional (i.e., effect-SNP) and those where it is not. For each factor motif, we then calculated the number of binding sites belonging to each of the four categories (divergent functional, divergent silent, polymorphic functional, and polymorphic silent) and calculated a selection score similar to the McDonald-Kreitman test (Supplemental S7.4).

### Expansion of GWAS Catalog

We created an expanded GWAS catalog by adding SNPs in linkage disequilibrium (LD) with each GWAS hit. Using 1000 Genomes data for European populations, we mapped each GWAS hit to all SNPs on the same LD block at an *r*^2^ cutoff of 0.8. The final file is a tabix-indexed bed file where each SNP entry has fields for its corresponding GWAS SNP, the *r*^2^ between them, and associated GWAS traits.

### Integrating functional annotations with GWAS

To integrate functional annotations and GWAS results, we used the fgwas command line tool (Pickrell 2014). fgwas computes association statistics genome wide using all common SNPs from European populations in the 1KG Project, splitting the genome into blocks larger than LD. Summary statistics were imputed with ImpG using *Z*-scores from meta-analysis data. Using an empirical Bayesian framework implemented in the fgwas software, GWAS data were then combined with functional annotations. We then compared the informativeness of these annotations from each of the 1891 motifs with CENTIPEDE predicted regulatory sites to a baseline model (see Supplemental S8.2) consisting of previously used genomic annotations identified as relevant (Pickrell 2014).

### Validating putative causal SNPs by reporter gene assays

We selected six of the 88 unique putatively causal GWAS SNPs. SNPs were chosen based on being in active footprints in LCLs (the cell line used for transfection), being effect-SNPs, and showing one or more of the following characteristics: allele specific binding (ASB), *PPA* > 0.5 in fgwas, or eQTL in the GTEX data. rs4519508 was chosen for having ASB; rs9686661, and rs6730558 were chosen for highest predictiveness; and rs2336384, rs532436, and rs4973431 for highest predictiveness and being an eQTL.

To test enhancer expression, we first constructed inserts containing the reference or alternate allele for each SNP of interest (see Supplemental S8.3). Cloning was performed using the Infusion Cloning HD kit (Clontech) and DNA was extracted using the PureYield kit (Promega). Transfections were performed into GM18507 using the standard protocol for the Nucleofector electroporation (Lonza). Luciferase activity was measured for four replicate experiments using the Dual-Glo Luciferase Assay Kit (Promega). We contrasted the activity of each construct to the pGL4.23 vector, to assess enhancer/repressor activity of each region. To evaluate allelespecific effects, we contrasted the activity of the reference allele to the alternate allele for each region and we used a t-test to assess significance.

## Data Access

Our footprint and footprint-SNP annotations are available as a custom UCSC Genome Browser hub. Please see http://genome.grid.wayne.edu/centisnps/ for information on how to view and download the data.

## Acknowledgements

Funding to support this research was provided by NIH 1 R01GM109215-01 (RPR and FL) and AHA14SDG20450118 (FL). All the computation was performed at the Wayne State University High Performance Computing Grid, and in the NSF supported Stampede Texas Advanced Computer Center (TACC) using an XSEDE start-up account (TG-MCB130080). We would like to thank Joe Pickrell for his assistance in running fgwas and in providing the GWAS meta-analysis datasets, Jacob Degner for reviewing an earlier version of this manuscript, and the members of the Luca/Pique group for helpful discussions.

